# Comprehensive Profiling of RBM20-Regulated CaMKIIδ Splice Variants Across the Heart, Skeletal Muscle, and Olfactory Bulbs

**DOI:** 10.1101/2025.02.27.640675

**Authors:** Yui Maeda, Yuri Yamasu, Hidehito Kuroyanagi

## Abstract

Calcium/calmodulin-dependent protein kinase IIδ (CaMKIIδ), encoded by the *Camk2d* gene, plays key regulatory roles in various Ca^2+^-regulated cellular processes. Extensive alternative splicing of the *Camk2d* gene generates multiple CaMKIIδ splice variants that exhibit differential roles. Despite significant advances in understanding the functions of CaMKIIδ, the full repertoire of *Camk2d* splice variants in a variety of tissues and their distinct roles in physiological and pathological contexts remain incompletely characterized due to the complex nature of multiple alternative splicing events in a single gene. Here, we conducted long-read amplicon sequencing to comprehensively investigate the murine *Camk2d* splice variants in the heart, skeletal muscle, and olfactory bulbs and show that *Camk2d* mRNAs in the heart and skeletal muscle have shorter 3’UTRs. We also demonstrate that a key regulator of *Camk2d* splicing, RNA-binding motif protein 20 (RBM20), whose *gain-of-function* mutations cause dilated cardiomyopathy, is crucial for the expression of heart-specific splice variants. Olfactory bulbs specifically express novel splice variants that utilize a mutually exclusive exon 6B and/or an alternative polyadenylation site in a novel exon 17.5. The tissue-specific repertoire of CaMKIIδ splice variants and their aberrant expression in disease model animals will help in understanding their roles in physiological and pathological contexts.

## Introduction

Calcium/calmodulin-dependent protein kinase II (CaMKII) is a serine/threonine kinase that plays a crucial regulatory role in various Ca^2+^-dependent cellular processes, including excitation-contraction coupling, synaptic plasticity, and gene transcription (Hudmon & Schulman, 2002). The CaMKII family is highly conserved among mammalian species and comprises four major isoforms (CaMKIIα, β, γ, and δ), each encoded by distinct genes and exhibiting tissue-specific expression patterns (Bayer & Schulman, 2019). Among these, CaMKIIδ, encoded by the *Camk2d* gene, is particularly abundant in cardiac and vascular tissues, where it plays a pivotal role in cardiovascular function and pathological remodeling (Anderson *et al*, 2011).

Alternative splicing of *Camk2d* generates multiple CaMKIIδ splice variants with distinct subcellular localization, substrate specificity, and functional properties (Duran *et al*, 2021; Gray & Heller Brown, 2014). In cardiac tissue, predominant splice variants include CaMKIIδ_1/A_,δ_2/C_,δ_3/B_, and δ_9_, each contributing uniquely to cardiac physiology and pathology. For instance, CaMKIIδ_1/A_ has been implicated in cardiac hypercontractility in conditional *Srsf1* knockout mice (Xu *et al*, 2005) and in isoproterenol-treated rats (Li *et al*, 2011a). CaMKIIδ_2/C_ has been associated with adverse effects in cardiomyocytes following *in vivo* ischemia/reperfusion injury (Gray *et al*, 2017) and pressure overload (Ljubojevic-Holzer *et al*, 2020). CaMKIIδ_3/B_ uniquely contains a nuclear localization signal (NLS) encoded by exon 14, leading to its predominant nuclear localization (Srinivasan *et al*, 1994). Transgenic overexpression of this cardiac-specific isoform induces cardiac hypertrophy (Zhang *et al*, 2002) but provides protection against ischemia/reperfusion injury (Gray *et al*., 2017). In contrast, cardiac-specific transgenic overexpression of CaMKIIδ_9_ results in cardiac hypertrophy, ventricular dilation, and cardiomyocyte death, ultimately progressing to severe heart failure (Zhang *et al*, 2019).

The alternative splicing of *Camk2d* is regulated *in vivo* by several RNA-binding proteins (RBPs), including serine/arginine-rich splicing factor 1 (SRSF1, also known as ASF/SF2) (Xu *et al*., 2005), RNA-binding motif protein 20 (RBM20) (Guo *et al*, 2012), RNA-binding Fox-1 homolog 2 (RBFOX2) (Wei *et al*, 2015), and RNA-binding motif protein 24 (RBM24) (Liu *et al*, 2022). Among these, RBM20 has garnered particular attention due to its strong association with an autosomal dominant form of familial dilated cardiomyopathy (DCM) (Brauch *et al*, 2009). Pathogenic *RBM20* mutations in familial DCM cases are highly enriched in a five-residue stretch RSRSP, which is essential for RBM20 nuclear localization (Murayama *et al*, 2018; Watanabe *et al*, 2018). These missense mutations function in a *gain-of-function* manner, as knock-in mice harboring patient-mimicking mutations exhibit DCM-like phenotypes, whereas knockout mice do not (Ihara *et al*, 2020; Methawasin *et al*, 2014; van den Hoogenhof *et al*, 2018). RNA sequencing (RNA-seq) analysis of human induced pluripotent stem cell-derived cardiomyocytes (iPSC-CMs) carrying an RSRSP motif mutation revealed aberrant alternative splicing patterns distinct from those observed in RBM20 knockout models (Fenix *et al*, 2021), suggesting a pathogenic role of splicing dysregulation in DCM. Recently, novel *RBM20* mutations located outside the RSRSP motif have been identified in familial DCM cases (Beqqali *et al*, 2016; Gaertner *et al*, 2020), but their effects on RBM20-mediated splicing regulation and DCM pathogenesis remain unexplored in animal models.

Beyond cardiac tissue, CaMKIIδ is also expressed in other tissues, including the brain (Sloutsky *et al*, 2020), skeletal muscle (Eigler *et al*, 2021), and vascular smooth muscle cells (Li *et al*, 2011b). Despite significant advances in understanding the function of CaMKIIδ, the full repertoire of *Camk2d* splice variants across various tissues and their distinct physiological and pathological roles remain incompletely characterized. In this study, we comprehensively investigate the diversity of *Camk2d* splice variants in murine heart, skeletal muscle, and brain. Furthermore, we assess the impact of an established DCM-associated *Rbm20* mutation (S637A) (Ihara *et al*., 2020) and a novel variant (V894A), which mimics the familial DCM mutation V914A (Gaertner *et al*., 2020), on *Camk2d* alternative splicing.

## Results

### *Rbm20* is expressed in ventricles, soleus, and olfactory bulbs

To determine whether the *Rbm20* gene is expressed in the soleus and olfactory bulbs in addition to the heart, we performed RT-PCR analysis using RNA extracted from these tissues. As our study focused on the effects of *Rbm20* mutations on *Camk2d* splicing, we utilized wild-type (*WT*), *Rbm20*^*KO/KO*^, *Rbm20*^*S637A/S637A*^, and *Rbm20*^*V894A/V894A*^ mice. RT-PCR revealed that *Rbm20* is expressed in both the soleus and olfactory bulbs, albeit at lower levels compared to the ventricles (Fig. 1).

**Figure 1.**
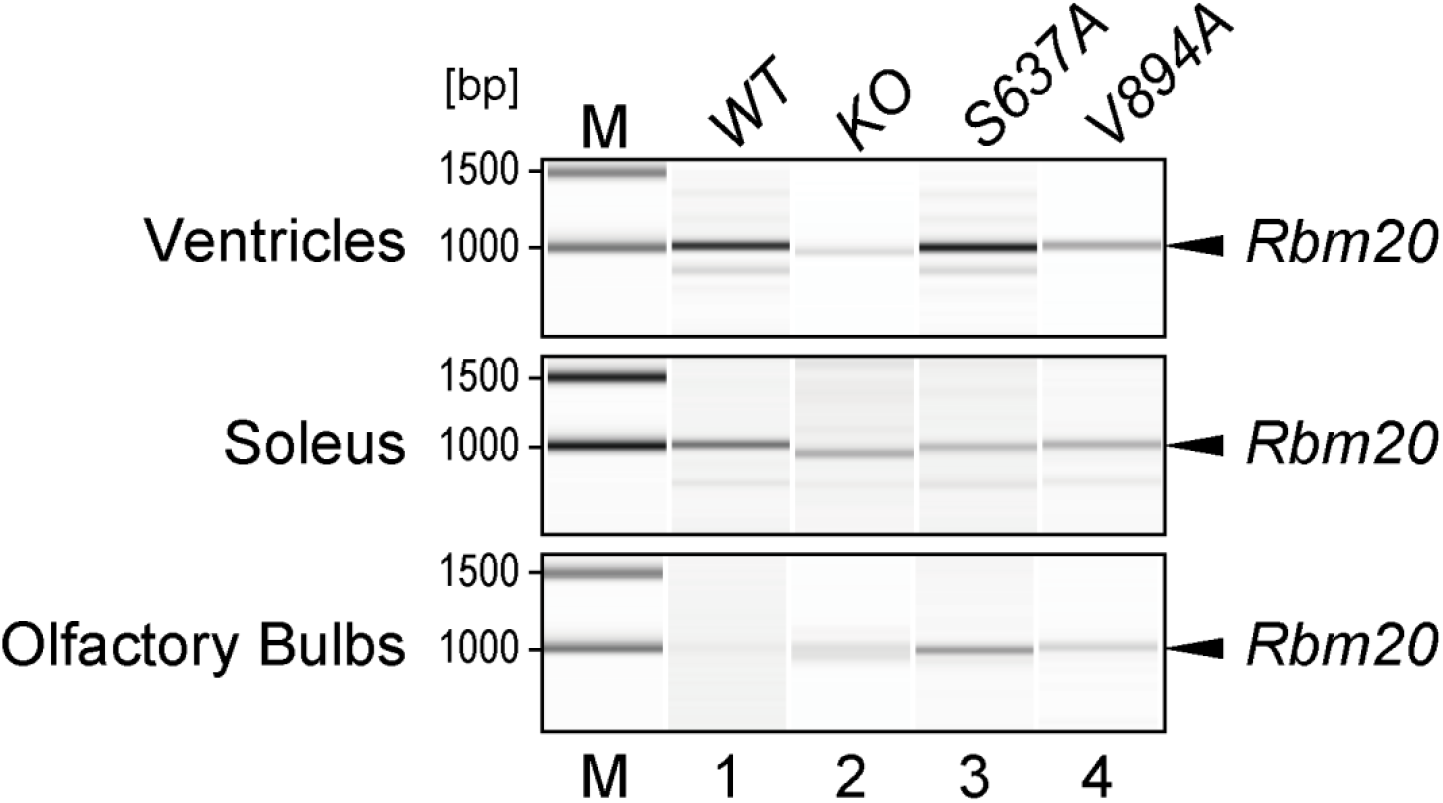
Expression of *Rbm20* in ventricles, soleus, and olfactory bulbs. RT-PCR analysis of *Rbm20* transcripts in ventricles (top), soleus (middle), and olfactory bulbs (bottom) from 8-to 9-week-old wild-type (lane 1), *Rbm20*^*KO/KO*^ (lane 2), *Rbm20*^*S637A/S637A*^ (lane 3), and *Rbm20*^*V894A/V894A*^ (lane 4) mice. Note that the RT-PCR products from *Rbm20*^*KO/KO*^ tissues (985 bp) were shorter than those from the other genotypes (1011 bp). M, DNA7500 ladder. Bioanalyzer-generated gel-like images (Agilent) are shown.

### Multiple *Camk2d* variants are expressed in ventricles, soleus, and olfactory bulbs

The Reference Sequence (RefSeq) database (release 227, NCBI) (O’Leary *et al*, 2016) lists nine splice variants of the murine *Camk2d* gene, including previously uncharacterized exons. The RefSeq sequences NM_001025439 and NM_001346636 contain an exon located downstream of, and homologous to, exon 6. Given that these exons were mutually exclusive in our experiments, we designated them as exons 6A and 6B (Fig. 2A). Additionally, NM_001293666 includes a novel long exon (2732 bp) situated between exons 17 and 18, which we termed exon 17.5. This exon contains a termination codon and five putative polyadenylation signals (PASs, AATAAA) (Fig. 2A). Exon 21, which also harbors a termination codon, is a cassette exon, and its skipping results in alternative C-terminal sequences for CaMKIIδ (Duran *et al*., 2021). The longest last exon (3569 bp) in other RefSeq sequences, exon 22, contains eight putative PASs (Fig. 2A), suggesting the presence of mRNAs with shorter 3’ untranslated regions (3’UTRs).

**Figure 2.**
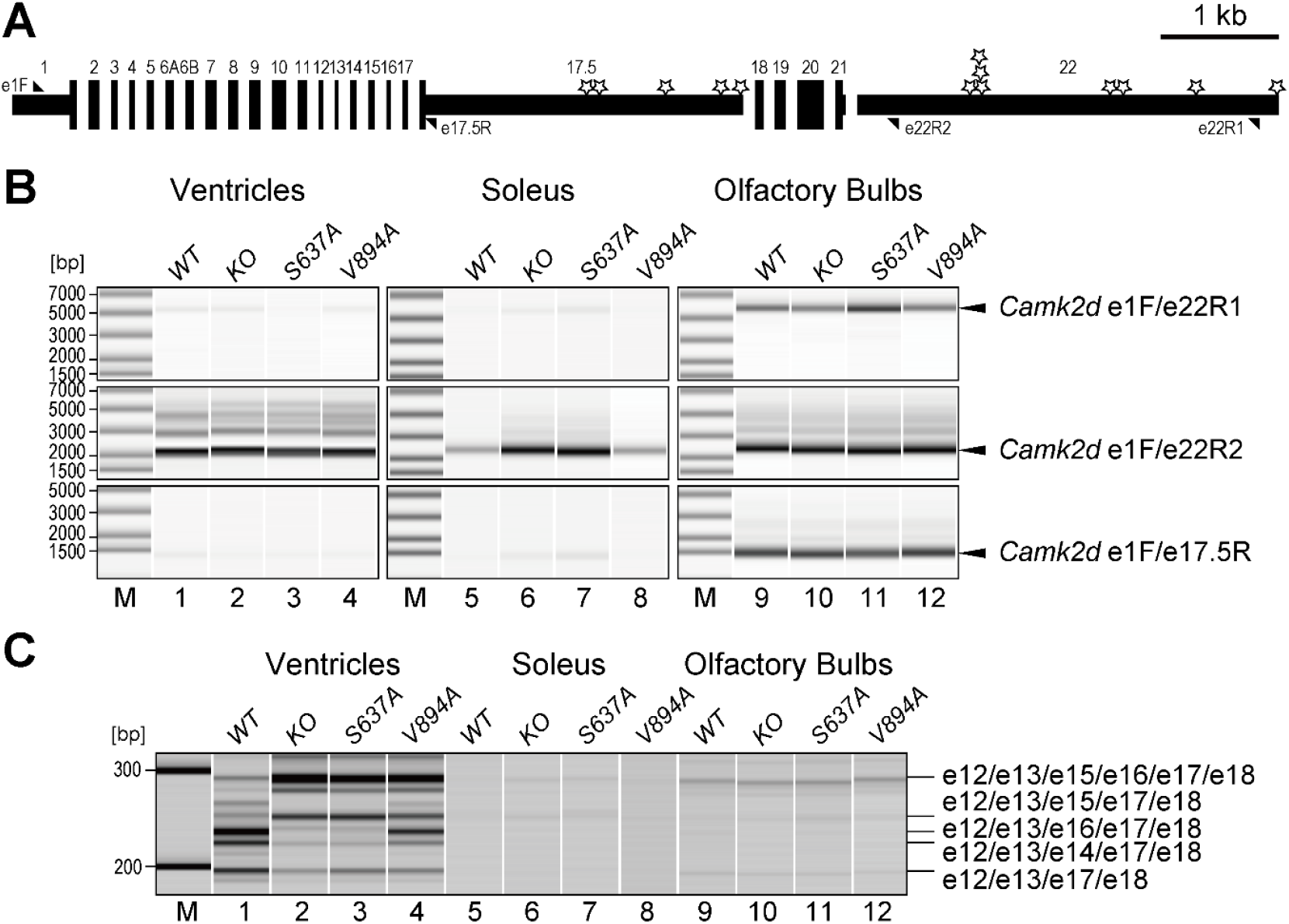
Expression of *Camk2d* isoforms in ventricles, soleus, and olfactory bulbs. (A) Schematic representation of the *Camk2d* gene structure. Exons are represented as boxes, with Untranslated regions (UTRs) depicted as thinner boxes. Triangles indicate primer positions. Stars denote putative PASs (AATAAA). Introns are not scaled. (B, C) RT-PCR analysis of *Camk2d* mRNAs in ventricles (lanes 1-4), soleus (lanes 5-8), and olfactory bulbs (lanes 9-12) from 8-to 9-week-old wild-type (lanes 1, 5, 9), *Rbm20*^*KO/KO*^ (lanes 2, 6, 10), *Rbm20*^*S637A/S637A*^ (lane 3, 7, 11), and *Rbm20*^*V894A/V894A*^ (lane 4, 8, 12) mice. Top: RT-PCR using primers e1F/e22R1 (longest 3’UTR). Middle: Primers e1F/e22R2 (all exon 22 isoforms). Bottom: Primers e1F/e17.5R (exon 17.5 isoforms). Bioanalyzer-generated gel-like images (Agilent) are shown. M, DNA ladder.

To verify the expression of these novel variants across tissues, we designed reverse primers targeting specific exon-polyadenylation site combinations. Primer e22R1 selectively amplified mRNAs with the longest 3’UTR, while e22R2 amplified all exon 22-containing mRNAs (Fig. 2A). We also designed e17.5R to detect mRNAs containing exon 17.5.

RT-PCR analysis revealed that *Camk2d* transcripts with the longest 3’UTR were readily detected in the olfactory bulbs but not in the ventricles or soleus (Fig. 2B, top panels), even though shorter 3’UTR-containing mRNAs were present in all tissues (Fig. 2B, middle panels). Similarly, mRNAs containing exon 17.5 were detected predominantly in the olfactory bulbs (Fig. 2B, bottom panels). Importantly, these isoforms were not significantly affected by any of the *Rbm20* mutations (Fig. 2B), suggesting that the selection among multiple PASs occurs in a tissue-specific, but RBM20-independent, manner.

To further investigate well-characterized *Camk2d* alternative splicing events involving exons 14-16, we performed RT-PCR using primers e12F and e18R. As expected, we confirmed RBM20-dependent regulation of these exons in the ventricles (Fig. 2C, lanes 1-4), consistent with previous findings (Ihara *et al*., 2020; Murayama *et al*., 2018). While signals were weaker in the soleus and olfactory bulbs due to lower RNA input, multiple RT-PCR products were detected in all tissues. Notably, the splicing patterns appeared unaffected in the *Rbm20* mutant mice (Fig. 2C, lanes 5-12). These results indicate that multiple *Camk2d* splice variants are expressed in a tissue-specific manner.

### Long-read sequencing reveals exon composition and expression levels of *Camk2d* splice variants

To comprehensively analyze the repertoire of *Camk2d* isoforms expressed in excitable tissues, we performed high-throughput long-read sequencing of full-length *Camk2d* amplicons obtained from the RT-PCR products in Figure 2B. Libraries were prepared by pooling three RT-PCR products per sample (amplified using e1F/e22R1, e1F/e22R2, and e1F/e17.5R) and barcoding them before sequencing. Twelve samples (from three tissues across four genotypes) were pooled and analyzed on a nanopore sequencing platform. Reads were mapped to classify transcripts based on 48 possible exon combinations.

Analysis of full-length sequencing reads from wild-type ventricles, soleus, and olfactory bulbs revealed tissue-specific exon selection (Fig. 3). The ventricles and soleus exclusively selected exon 6A, whereas olfactory bulbs expressed a subset of isoforms containing exon 6B. Alternative splicing patterns of the CaMKIIδ variable linker domain (exons 14-16) also varied across tissues. In ventricles, predominant isoforms included e16-only variants (δ_9_,δ_10_), e14-only variants (δ_3/B_,δ_7_), and exon 14/15/16-skipped variants (δ_2/C_,δ_6_). These findings are consistent with prior long-read amplicon sequencing of murine *Camk2d* transcripts (Zhang *et al*., 2019), validating our methodology.

**Figure 3.**
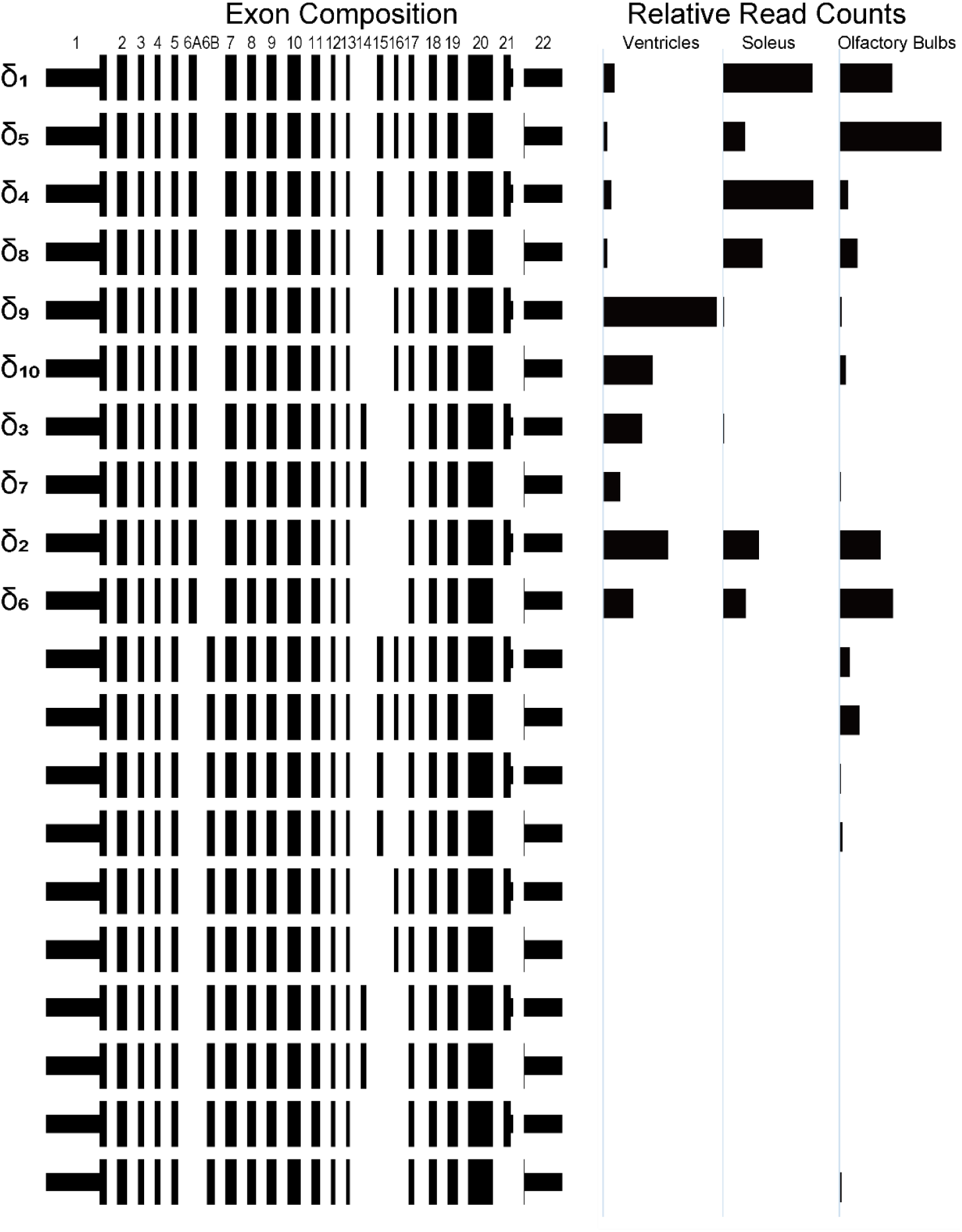
Long-read sequencing of *Camk2d* splice variants in a wild-type mouse. Left: Schematic representation of *Camk2d* splice variants. Right: Relative read counts of cDNA amplicons (e1F/e22R2) from wild-type ventricles (1641 reads), soleus (360 reads), and olfactory bulbs (2615 reads). Variants are categorized based on exon 6A/6B selection. Isoforms with frequencies <1% were omitted.

### RBM20 regulates heart-specific alternative splicing of the *Camk2d* gene

To determine the impact of *Rbm20* mutations on *Camk2d* alternative splicing, we analyzed transcript isoforms in ventricles, soleus, and olfactory bulbs from *Rbm20*^*KO/KO*^, *Rbm20*^*S637A/S637A*^, and *Rbm20*^*V894A/V894A*^ mice (Fig. 4).

**Figure 4.**
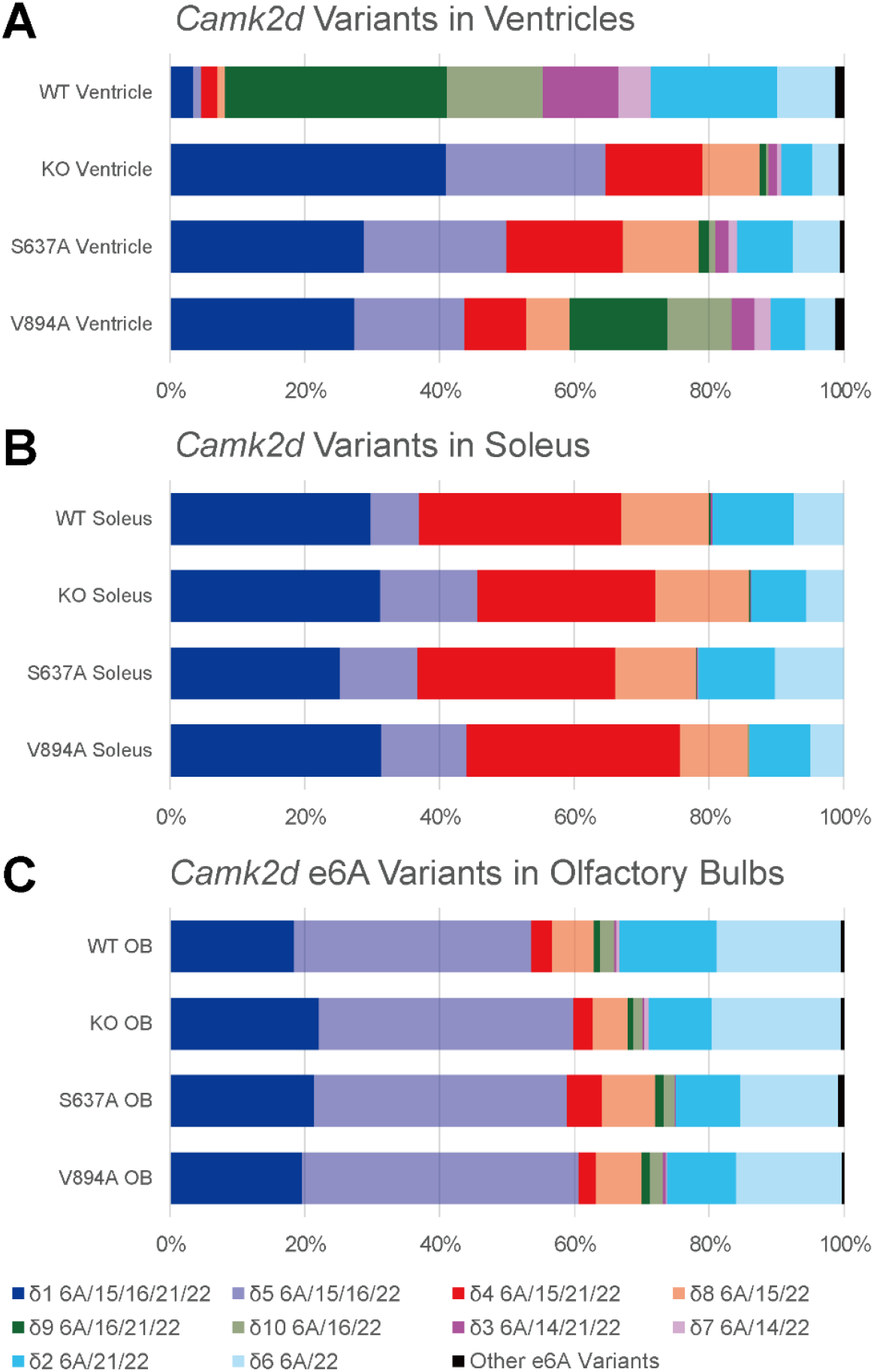
RBM20 regulates heart-specific alternative splicing of *Camk2d*. (A-C) Proportions of *Camk2d* splice variants in ventricles (A), soleus (B), and olfactory bulbs (C) from 8-to 9-week-old animals of the indicated genotypes. Only variants containing exon 6A are shown; other isoforms specific to the olfactory bulbs are analyzed in Figure 5.

As expected, *Rbm20* knockout and the S637A mutation drastically altered the proportions of *Camk2d* splice variants in the ventricles (Fig. 4A), consistent with the RT-PCR results (Fig. 2C). Notably, the ratio of exon 21-included to exon 21-skipped isoforms remained unaffected by *Rbm20* mutations, suggesting that exon 21 regulation is independent of RBM20 activity (Fig. 4A). The *Rbm20*^*V894A*^ mutation exhibited a modest effect on RBM20-mediated alternative splicing, indicating a partial *loss-of-function* phenotype for *Camk2d* regulation.

In contrast, *Camk2d* isoform proportions in the soleus (Fig. 4B) and olfactory bulbs (Fig. 4C) remained largely unaffected by *Rbm20* mutations, despite detectable *Rbm20* expression in these tissues (Fig. 1). These findings confirm that RBM20-dependent regulation is specific to the heart, influencing the expression of ventricular-enriched *Camk2d* splice variants, including δ_9_,δ_10_,δ_3_/_B_, and δ_7_.

### Olfactory bulbs express unique splice variants of the *Camk2d* gene

To further characterize the unique repertoire of *Camk2d* transcripts in olfactory bulbs, we performed an additional round of long-read amplicon sequencing using RNA from wild-type and *Rbm20* mutant mice (Fig. 5).

**Figure 5.**
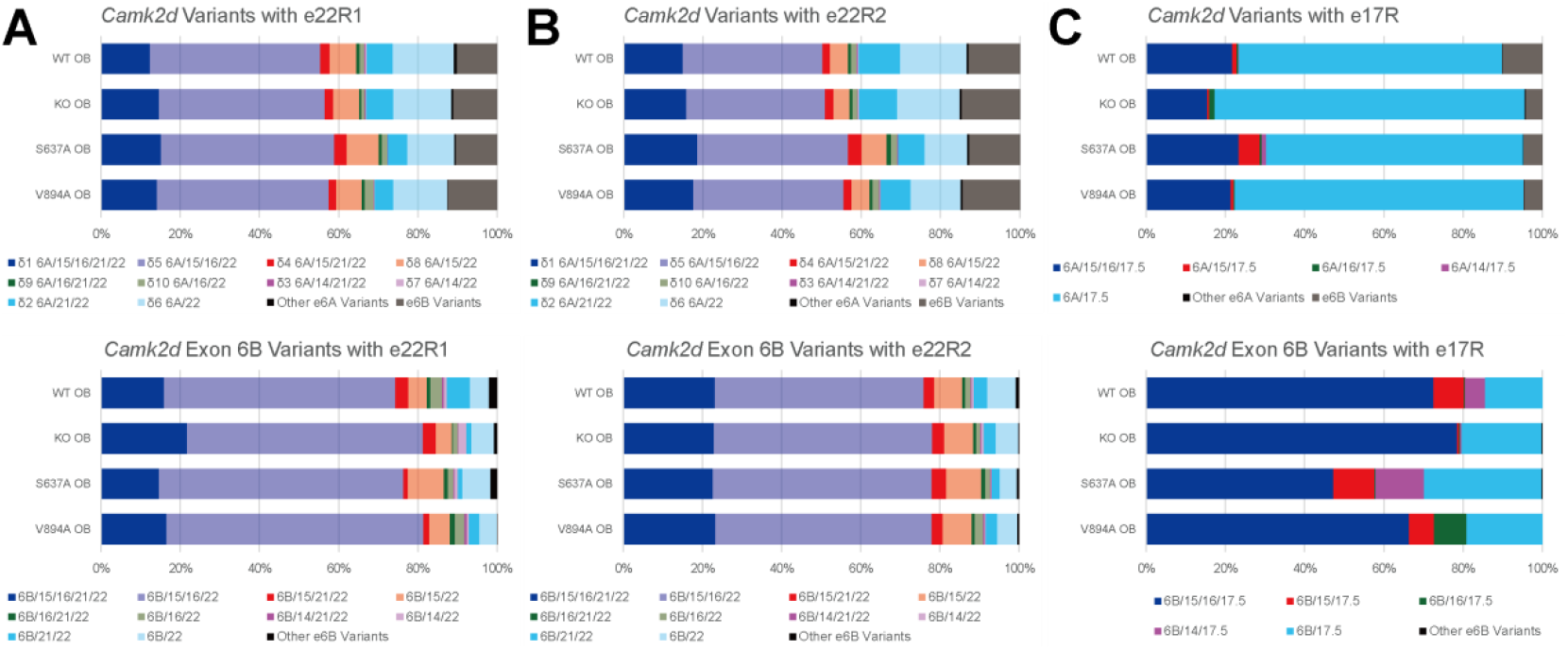
Unique splice variants of *Camk2d* expressed in olfactory bulbs. (A-C) Proportions of *Camk2d* splice variants expressed in olfactory bulbs, amplified using reverse primers e22R1 (A), e22R2 (B), and e17.5R (C) from 8-to 9-week-old animals of the indicated genotypes. Top panels: proportions of all detected variants. Bottom panels: proportions of exon 6B-containing variants (gray in Top panels).

We obtained full-length sequencing reads from amplicons amplified using e22R1, which selectively targets exon 22-containing transcripts with the longest 3’UTR (Fig. 2A). Among these, exon 6B-containing variants accounted for 10.2%–12.4% of total reads, indicating alternative exon 6 selection in a subset of olfactory bulb *Camk2d* mRNAs (Fig. 5A, top panel). Alternative splicing patterns of exons 14-16 in these exon 6B variants were similar to those observed for exon 6A variants, though inclusion of exons 15 and 16 was more prevalent (Fig. 5A, bottom panel).

To determine whether exon 22 splice variant distributions differed based on 3’UTR length, we analyzed amplicons amplified with e22R2, which detects all exon 22-containing mRNAs. The proportions of splice variants detected using e22R2 (Fig. 5B) were highly similar to those obtained using e22R1 (Fig. 5A), suggesting that most, if not all, exon 22-containing mRNAs in olfactory bulbs possess the longest 3’UTR.

Additionally, sequencing of exon 17.5-containing variants (amplified using e17.5R) revealed unique alternative splicing patterns in olfactory bulbs. The majority of *Camk2d* mRNAs containing exon 17.5 lacked exons 14, 15, and 16 (Fig. 5C). Notably, 4.2%–10.0% of exon 17.5-containing transcripts included exon 6B, with most of these also retaining exons 15 and 16 (Fig. 5C).

Importantly, *Rbm20* mutations did not significantly alter the proportions of *Camk2d* transcripts in olfactory bulbs (Fig. 5A-C), reinforcing the conclusion that olfactory bulb-specific alternative splicing is independent of RBM20. These results indicate that the olfactory bulbs express a unique set of *Camk2d* isoforms, including mRNAs with the longest 3’UTR, exon 6B variants, and exon 17.5-containing transcripts with distinct C-terminal and 3’UTR structures, none of which are regulated by RBM20.

## Discussion

In this study, we conducted long-read amplicon sequencing of *Camk2d* transcripts and demonstrated several key findings: (i) the olfactory bulbs predominantly express mRNAs with the longest 3’UTR, whereas the ventricles and soleus preferentially express mRNAs with shorter 3’UTRs (Fig. 2B); (ii) wild-type ventricles express ten major *Camk2d* splice variants containing exon 6A, consistent with previous reports (Zhang *et al*., 2019), whereas the soleus expresses only six of these isoforms (Figs. 3 and 4A); (iii) expression of four ventricular-specific isoforms is absolutely dependent on RBM20 (Fig. 4A); (iv) the olfactory bulbs uniquely express novel *Camk2d* variants containing exons 6B and/or 17.5, in addition to those containing exons 6A and 22 (Figs. 3 and 5); and (v) despite detectable *Rbm20* expression in the soleus and olfactory bulbs (Fig. 1), the proportions of *Camk2d* splice variants in these tissues remain largely unaffected by *Rbm20* mutations (Figs. 4 and 5).

Long-read sequencing provided a comprehensive view of *Camk2d* alternative splicing and polyadenylation, revealing distinct selection patterns for each event. Exon 14, which encodes an NLS (Srinivasan *et al*., 1994), is included exclusively in the ventricles in an RBM20-dependent manner (Figs. 3 and 4). Notably, exon 14 is never co-included with exon 15 or 16, regardless of tissue type or *Rbm20* genotype (Figs. 3-5). Exon 21 inclusion and exclusion are observed across all examined tissues (Fig. 3). Within each tissue, the ratio of exon 21-included to exon 21-excluded variants remains consistent among transcripts with otherwise identical splicing patterns, although this ratio differs among tissues (Fig. 4). This observation suggests that exon 21 regulation occurs in a tissue-specific manner, independent of other alternative exons.

Exon 6B is utilized in approximately 10% of *Camk2d* transcripts exclusively in the olfactory bulbs (Fig. 5). The presence of exon 6B has also been predicted in the human *CAMK2D* gene based on RefSeq models (O’Leary *et al*., 2016) and short-read amplicon sequencing (Sloutsky *et al*., 2020). The amino acid sequences encoded by exons 6A and 6B are highly conserved between humans and mice (Fig. 6), suggesting a conserved functional role. The physiological significance of tissue-specific regulation of these mutually exclusive exons remains to be elucidated. The alternative exon 17.5 is also specifically utilized in the olfactory bulbs (Fig. 2). Splicing patterns of other *Camk2d* exons in exon 17.5-containing transcripts differ from those in exon 22-containing transcripts (Fig. 5), suggesting coordinated regulation of upstream exons and alternative polyadenylation at exon 17.5, likely in a neuron-type-specific manner. Given that longer 3’UTRs generated by alternative polyadenylation frequently serve as binding sites for microRNAs (Pereira-Castro & Moreira, 2021), *Camk2d* mRNAs in the olfactory bulbs—characterized by the longest 3’UTRs—may be subjected to an additional layer of post-transcriptional regulation via translational control.

**Figure 6.**
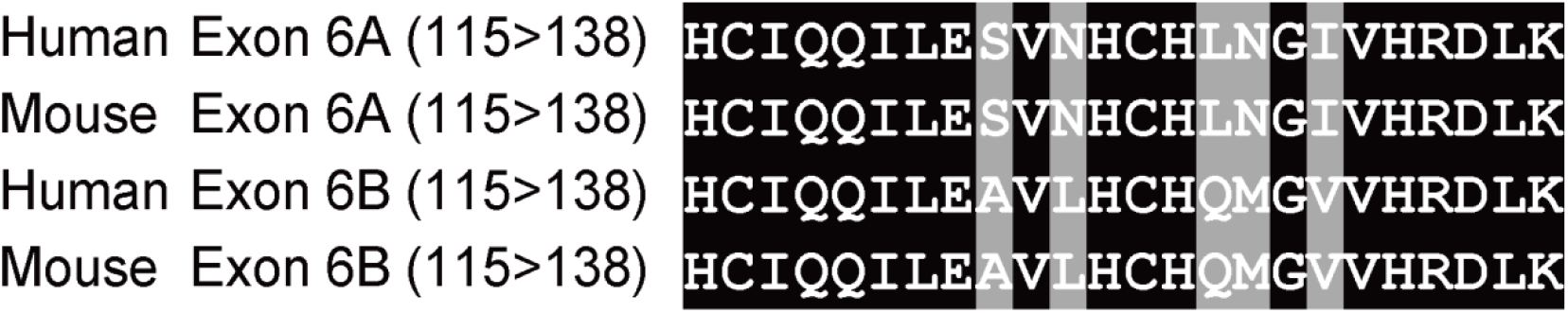
Evolutionary conservation of CaMKIIδ exon 6A/6B sequences. Amino acid sequence alignment of the CaMKIIδ protein kinase domain encoded by exons 6A and 6B from human and mouse. Identical residues across all sequences are highlighted in black, while residues differing between exons 6A and 6B are shaded in gray. Amino acid positions in each protein are indicated.

*Gain-of-function* mutations within the RSRSP stretch of *RBM20*, which result in the formation of aberrant cytoplasmic RBM20 condensates, have been strongly associated with DCM in human patients (Schneider *et al*, 2020) and animal models (Ihara *et al*., 2020; Kornienko *et al*, 2023; Nishiyama *et al*, 2022; Schneider *et al*., 2020; Wang *et al*, 2022). In this study, we also analyzed the effect of the *Rbm20*^*V894A*^ mutation (Y.Y. and H.K., unpublished) on *Camk2d* splicing and demonstrated that this mutation reduces RBM20 function, at least in terms of *Camk2d* alternative splicing (Fig. 4A). Among the RBM20-regulated genes, aberrant *Camk2d* splicing has been proposed as a key contributor to DCM-like phenotypes (Lennermann *et al*, 2020). However, it remains unclear whether mis-splicing of *Camk2d* or other RBM20 target genes directly contributes to DCM pathology, as altered splicing coincides with the formation of cytoplasmic RBM20 condensates. Notably, pathogenic *RBM20* mutations in human iPSC-CMs (Fenix *et al*., 2021; Nishiyama *et al*., 2022; Wyles *et al*, 2016; Zhu *et al*, 2021) and a pig model (Schneider *et al*., 2020) cause widespread alterations in the splicing of over 100 genes associated with cardiac function. Despite these widespread splicing changes, *Camk2d* is one of the few genes consistently affected in murine *Rbm20* knock-in and knockout models (Ihara *et al*., 2020; Yamamoto *et al*, 2022).

The detailed analysis of full-length *Camk2d* mRNA isoforms presented in this study provides valuable insights into the role of RBM20 in cardiac splicing regulation. These findings will contribute to future investigations into how pathogenic *Rbm20* mutations affect cardiomyocyte function and potentially influence alternative splicing in other tissues.

### Experimental Procedures

#### Animals

All animal experiments were conducted in accordance with the *Guide for the Care and Use of Laboratory Animals* (National Research Council, The National Academies Press, 8th edition, 2011). The study protocol was approved by the Institutional Animal Care and Use Committee of the University of the Ryukyus (Approval Nos. A2021026 and A2024042).

The Generation of the *Rbm20*^*S637A*^ knock-in allele (Murayama *et al*., 2018) and the *Rbm20*^*KO*^ allele (Ihara *et al*., 2020) has been described previously. The generation of the *Rbm20*^*V894A*^ knock-in allele will be reported elsewhere (Y.Y. and H.K., unpublished). Briefly, this allele was introduced using a cloning-free CRISPR/Cas system (Aida *et al*, 2015). A wild-type (*WT*) mouse strain C57BL/6J was used as controls.

#### RNA extraction and RT-PCR

Total RNA was extracted from heart ventricular tissues of anesthetized 8-to 9-week-old male mice using the RNeasy Mini Kit (QIAGEN) with on-column DNase I treatment (QIAGEN), following the manufacturer’s protocol for fibrous tissues. Complementary DNA (cDNA) was synthesized from total RNA using the PrimeScript II 1st strand cDNA Synthesis Kit (Takara). The RNA input amounts were as follows: 1.6 μg for ventricles, 0.32 μg for soleus muscle, and 0.24 μg for olfactory bulbs.

Semi-quantitative RT-PCR was performed to analyze alternative splicing patterns using PrimeSTAR GXL DNA Polymerase (Takara) or KOD One PCR Master Mix (TOYOBO). PCR primers were synthesized by Eurofins or Merck, with sequences provided in Supplementary Table S1. PCR products were analyzed using a Bioanalyzer 2100 Expert system with DNA7500 or DNA1000 Kits (Agilent).

**Table S1.**
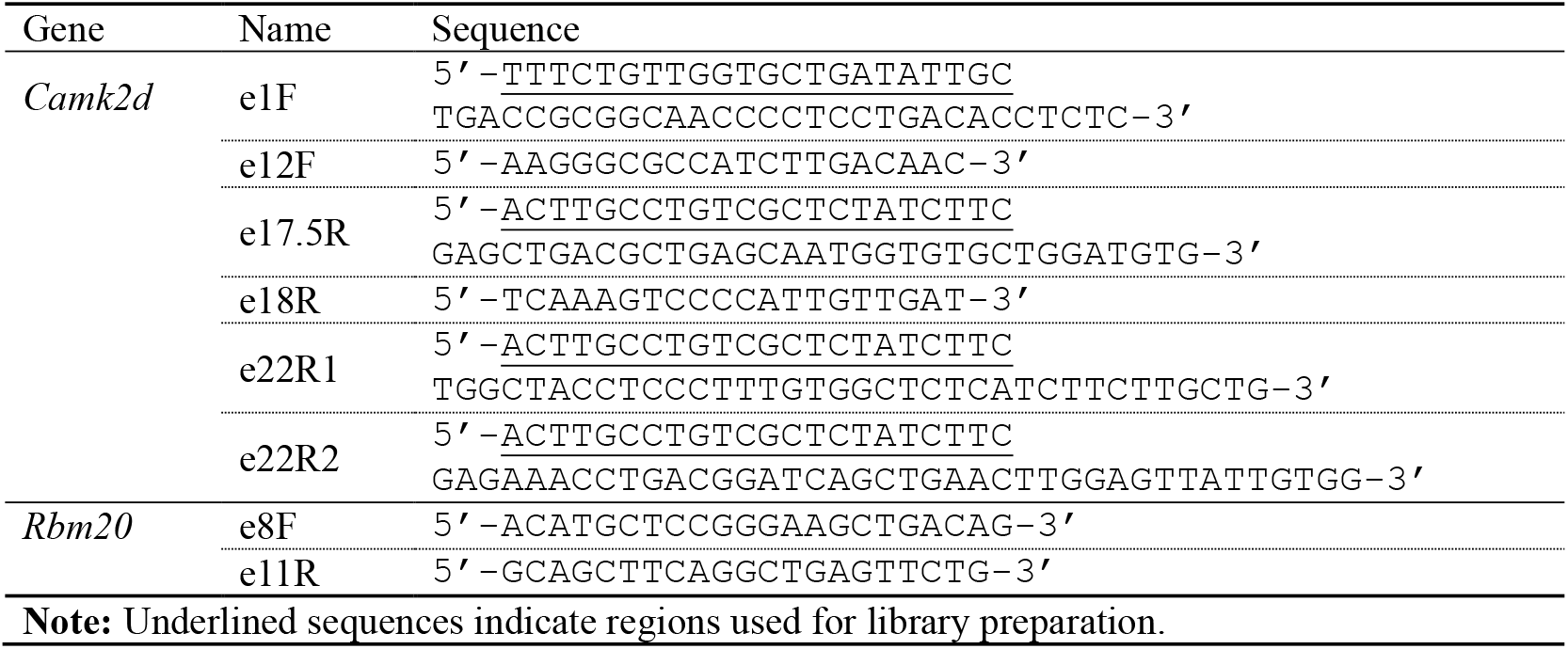
Primer Sequences Used in RT-PCR Assays and Library Preparation.

### Library preparation and long-read sequencing

Long-range PCR for amplicon sequencing was performed using KOD One PCR Master Mix (TOYOBO) following the manufacturer’s step-down protocol. Primer sequences are listed in Supplementary Table S1.

Nanopore amplicon sequencing libraries were prepared using the Ligation Sequencing Kit V14 (SQK-LSK114) and PCR Barcoding Expansion Pack 1-12 (EXP-PBC001) (Oxford Nanopore Technologies), according to the manufacturer’s instructions. Long-read sequencing was performed on a MinION Mk1B platform (Oxford Nanopore Technologies) using Flongle or MinION/GridION Flow Cells, following the standard protocol.

### Data analysis

Raw sequencing reads were processed using default parameters. Transcript isoform analysis was performed using IsoQuant 3.6 (Prjibelski *et al*, 2023), with *Mus musculus* reference genome data (mm39) obtained from the UCSC Genome Browser (Perez *et al*, 2025). A custom-prepared annotation file in GTF format was used for mapping *Camk2d* transcripts.

Amino acid sequence alignments of the CaMKIIδ protein kinase domain were conducted using the MegAlign Pro module of Lasergene Molecular Biology Ver. 18 (DNASTAR).

## Declarations

### Data availability

Long-read sequencing data will be available on a public database and is available upon request to H.K.

### Author Contributions

Y.M., Y.Y., and H.K. contributed to molecular and animal experiments. Y.Y. and H.K. contributed to data analysis. H.K. wrote the manuscript.

## Acknowledgements

We thank Akiko ISHII and Kuniko OSHIRO for their technical assistance in the maintenance of the gene-modified mice.

